# Myeloid cell and transcriptome signatures associated with inflammation resolution in a model of self-limiting acute brain inflammation

**DOI:** 10.1101/486589

**Authors:** Claire L Davies, Anirudh Patir, Barry W McColl

**Author notes:** **Correspondence:** Dr Barry McColl.

## Abstract

Inflammation contributes to tissue repair and restoration of function after infection or injury. However, some forms of inflammation can cause tissue damage and disease, particularly if inappropriately activated, excessive, or not resolved adequately. The mechanisms that prevent excessive or chronic inflammation are therefore important to understand. This is particularly important in the central nervous system where some effects of inflammation can have particularly harmful consequences, including irreversible damage. An increasing number of neurological disorders, both acute and chronic, and their complications are associated with aberrant neuroinflammatory activity.

Here we describe a model of self-limiting acute brain inflammation optimised to study mechanisms underlying inflammation resolution. Inflammation was induced by intracerebral injection of lipopolysaccharide (LPS) and the temporal profile of key cellular and molecular changes were defined during the progression of the inflammatory response. The kinetics of accumulation and loss of neutrophils in the brain enabled well demarcated phases of inflammation to be operatively defined, including induction and resolution phases. Microglial reactivity and accumulation of monocyte-derived macrophages were maximal at the onset of and during the resolution phase. We profiled the transcriptome-wide gene expression changes at representative induction and resolution timepoints and used gene coexpression network analysis to identify gene clusters. This revealed a distinct cluster of genes associated with inflammation resolution that were induced selectively or maximally during this phase and indicated an active programming of gene expression that may drive resolution as has been described in other tissues. Induction of gene networks involved in lysosomal function, lipid metabolism and a comparative switch to MHC-II antigen presentation (relative to MHC-I during induction) were prominent during the resolution phase. The restoration and/or further induction of microglial homeostatic signature genes was notable during the resolution phase.

We propose the current model as a tractable reductionist system to complement more complex models for further understanding how inflammation resolution in the brain is regulated and as a platform for *in vivo* testing/screening of candidate resolution-modifying interventions. Our data highlight how resolution involves active cellular and transcriptome reprogramming and identify candidate gene networks associated with resolution-phase adaptations that warrant further study.

## Introduction

Acute inflammation is critical to stimulate tissue repair yet if uncontrolled or insufficiently resolved can progress to chronic inflammation that has potentially harmful consequences^1^. In the central nervous system (CNS), chronic neuroinflammation and dysfunction of cells involved (e.g. microglia, recruited immune cells) is increasingly implicated in the progression of neurodegenerative disease ^2–6^. Infection as well as sterile damage to the CNS such as caused by stroke and trauma, also elicit potent inflammatory responses^7,8^. Chronic inflammatory responses to these acute events have been proposed as a potential cause of long-term degenerative and cognitive complications^9–11^. There is therefore a need to better understand mechanisms and to develop strategies that prevent progression to harmful chronic neuroinflammation. Understanding how self-limiting acute inflammatory reactions in the CNS are regulated may help achieve this.

Acute CNS inflammation is underpinned by proliferation and activation of brain-resident microglia^12^ and, in some cases, influx of blood-borne immune cells, notably neutrophils^13,14^ and monocyte-derived macrophages^15–17^. Collectively, these myeloid cells are key orchestrators of the innate immune response to CNS damage. Beneficial functions of inflammatory cells include phagocytosis of debris ^18–20^, provision of CNS barrier support ^21,22^, and trophic support for tissue healing and repair ^23–26^. Importantly, depletion of certain innate immune cells can exacerbate CNS injury and prevent timely repair ^25,27–29^. However, in some circumstances inflammation can be toxic and maladaptive. Following cerebral ischemia, failure to resolve the initial inflammatory response by removal of neutrophil toxins and apoptotic cells can exacerbate injury ^30^. The impact of controlling the severity of inflammation has been investigated through immunomodulatory interventions. For example, blocking interleukin-1, a critical mediator driving innate neuroinflammation ^31^, improved outcome in experimental stroke models ^32^ and has shown promise in clinical trials ^33,34^ Minocycline was promising in clinical trials following ischaemic stroke ^35–37^. Nonetheless, other anti-inflammatory treatments have been less successful in stroke^38,39^. In traumatic brain injury, treatment with methylprednisolone had a less favourable recovery ^40^ whilst progesterone had no effect ^41^. These variable outcomes highlight that broad suppression of inflammatory responses may also negate possible beneficial effects of inflammation required for tissue repair, and that a greater understanding of the endogenous mechanisms that drive favourable resolution in the CNS is warranted.

Inflammation resolution in tissues outside the CNS has been more extensively studied, often using reductionist models such as LPS or zymosan-induced peritonitis^42^. The utility of such models is underscored by their revealing insight to fundamental principles including that resolution is a highly active process and their identification of new mediators, notably the specialised pro-resolving mediators (SPMs) such as lipoxins, resolvins and protectins ^42–45^. Previous work highlighted that inflammatory mechanisms in the brain may differ from peripheral tissues ^31^ therefore it is unclear whether similar mechanisms and mediators operate during resolution of CNS inflammation. A tractable model of CNS inflammation optimised for probing cellular and molecular components of the resolution phase is likely to be a useful complement to more complex injury- and disease-specific models.

Here, we present a self-limiting model of acute CNS inflammation induced by intracerebral liposaccharide injection that has a well-demarcated resolution phase. We have determined the cellular profile and kinetics of key myeloid cell accumulation as the inflammatory reaction progresses from induction to resolution. We have also generated a transcriptomic profile defining resolution that highlights the active nature of the resolution phase and identifies candidate pathways and mediators for future study. The current work provides an experimental platform and underpinning data to support further studies addressing candidate endogenous mechanisms of inflammation resolution in the brain.

## Materials and Methods

### Mice

Male 8-12 week old C57b1/6J mice were purchased from Charles River Laboratories and acclimatised for a minimum of one week prior to procedures. A breeding colony of *Ccr2*^RFP^ mice (B6.129(Cg)-*Ccr2*^*tm2.1Ifc*^/J; Jax stock #017586) as described previously^46^ was maintained locally. Mice were maintained under specific pathogen-free conditions and a standard 12 h light/ dark cycle with unrestricted access to food and water. Mice were housed in individually ventilated cages in groups of up to five mice. All procedures involving live animals were carried out under the authority of a UK Home Office Project Licence in accordance with the ‘Animals (Scientific Procedures) Act 1986’ and Directive 2010/63/EU and were approved by the local Animal Welfare and Ethics Review Board (AWERB).

### Intracerebral injections

Mice were positioned in a stereotaxic frame (Stoelting, USA) under isoflurane anaesthesia (0.2 L/min O_2_ and 0.5 L/min N_2_O). The skull was exposed via a midline incision and a small craniotomy was made overlying the left hemisphere 2 mm lateral to Bregma using an Ideal micro drill (tip diameter 0.8 mm) (Stoelting). Stereotaxic injections were performed using pre-calibrated glass microcapillary pipettes (Drummond Scientific, USA), previously pulled using a vertical electrode puller (Model PP830, Japan). Co-ordinates for intrastriatal injection were 2 mm lateral to Bregma (left hemisphere) and 2.5 mm below the brain surface (Franklin and Paxinos, 2007). 1 μL LPS (*E. coli* 0127:B8, 5 mg/ml) or PBS was injected at a rate of 0.5 μL/min. The wound was sutured and topical local anaesthetic (lidocaine/prilocaine, EMLA) applied. Mice were recovered from general anaesthesia and returned to standard housing for indicated durations. Post-operative dehydration was prevented with subcutaneous saline injections. Mice were monitored regularly during the post-operative period using a clinical scoring system.

### Tissue processing

At indicated time points, mice were deeply anaesthetised with isoflurane (0.2 L/min O_2_ and 0.5 L/min N_2_O) and perfused transcardially with physiological saline (0.1 % DEPC-treated for experiments proceeding to RNA extraction) to remove circulating blood. For flow cytometry, the hemisphere ipsilateral to injection was dissected and a brain cell suspension prepared immediately as below. For transcriptomic profiling, a 4 mm brain sample block surrounding the injection site was dissected from the ipsilateral hemisphere using a mouse brain matrix and snap-frozen. For immunohistochemistry and histology, after saline perfusion mice were perfused with 4% paraformaldehyde. Brains were removed and post-fixed in 4% paraformaldehyde for 24-48 h, cryoprotected in 20% sucrose solution overnight then frozen by immersion in chilled (−40°C) isopentane. 20 μm coronal sections were cut on a freezing sledge microtome (Bright Instruments, Huntingdon, UK) and stored in cryoprotectant medium (100 ml 10x phosphate buffer, 300 ml ethylene glycol, 200 ml glycerol, 400 ml dH_2_O) at −20°C.

### Immunohistochemistry

Cryoprotected brain sections were placed into a 24-well plate and washed in PBS. Endogenous peroxidase was blocked by incubation with 0.3% hydrogen peroxide in dH_2_O for 15 min and sections washed in PBS. Sections were incubated in 5% normal serum (from species secondary antibody raised in) diluted in 0.3% Triton/PBS for 1 h then in primary antibody solution (2.5% normal serum in 0.3% Triton/PBS) overnight at 4°C. Primary antibodies used were: rabbit polyclonal anti-IBA1 (Wako Chemicals, Japan), rabbit polyclonal anti-neutrophils (batch SJC4, gift from D Anthony, University of Oxford, UK), goat polyclonal anti-ICAM-1 (R&D Systems, UK). Primary antibody staining was omitted for detection of endogenous mouse IgG. Sections were then washed in PBS and incubated with biotinylated secondary antibody (1:200 in PBS; Vector Laboratories, UK) for 1 h. After further washes in PBS, sections were incubated with avidin-biotin peroxidase complex solution (Vectastain ABC, Vector, UK) for 1 h, washed in PBS then incubated in diaminobenzidine (DAB) solution (100 μl DAB, 2.5 μl H_2_O_2_ in 5 ml dH_2_0) for 30 s. DAB solution was aspirated and sections rinsed with dH_2_O. Sections were washed in PBS, mounted on gelatine-coated slides and dried overnight then dehydrated in a series of alcohols followed by incubation in xylene for 10 min. Coverslips were applied using Pertex medium (Cell Path Ltd., Newtown Powys, UK).

To quantify neutrophil accumulation on SJC4-immunostained sections, images were captured using a Nikon N1 upright microscope (Nikon, Surrey, UK). Regions-of-interest (ROI) were defined according to neuroanatomical reference points around the site of injection (**Figure 1A**). SJC4^+^ cells were marked on the digital images In Image J using the cell counter to ensure they were only counted once. The number of SJC4^+^ cells was counted in each of the ROIs and data expressed as the sum of the numbers in all ROIs.

**Figure 1.**
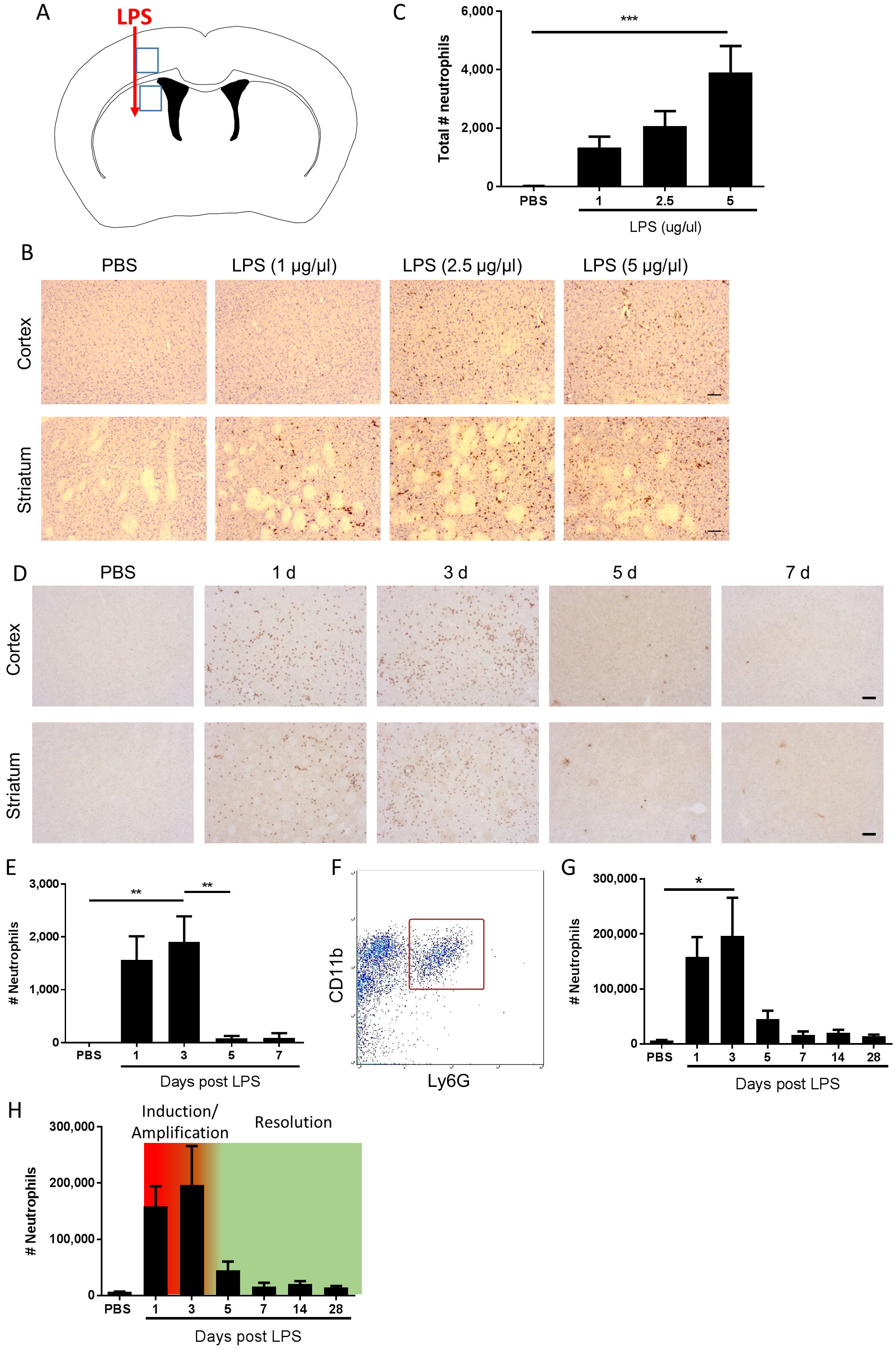
Kinetics of neutrophil accumulation and loss define operational phases of the inflammatory response to intracerebral LPS challenge. (A) Schematic showing location of stereotaxic injections. Regions 1 and 2 indicate areas of cerebral cortex and striatum, respectively, shown in B and D. (B) Representative immunostaining images of SJC4+ neutrophil accumulation in cortex (top) and striatum (bottom) (regions indicated in A) 24 h after LPS injection. Scale bar, 100 μm. (C) Quantification of the total number of SJC4+ neutrophils in the ipsilateral hemisphere 24 h after PBS or LPS injection. ***P < 0.001, one-way ANOVA with Dunnett’s post-hoc test. N = 4-5 mice/group. (D) Representative immunostaining images of SJC4+ neutrophils in the cortex (top) and striatum (bottom) (regions indicated in A) following PBS or LPS injection and mice recovered up to 7 d. (E) Quantification of the total number of SJC4+ neutrophils following PBS or LPS injection and recovered up to 7 d. **P < 0.01, one-way ANOVA with Bonferroni’s post-hoc test. N = 6 mice/group. (F) Flow cytometric identification of CD11b^+^ Ly6G^+^ neutrophils in brain cell suspensions. (G) Quantification of total number of CD11b^+^ Ly6G^+^ neutrophils following PBS or LPS injection and mice recovered for up to 28 d. *P < 0.05, one-way ANOVA with Bonferroni’s post-hoc test. N = 4-7 mice/group. (H) Kinetics of neutrophil accumulation and loss following LPS injection shown in G overlaid with the timings for the phases defined operatively in the current study as inflammation initiation/amplification (1-3 d) and resolution (3-5 d).

### Histology

Cryoprotected brain sections were washed in PBS, mounted on to gelatine-coated slides and dried overnight at room temperature. Sections were submerged in xylene, followed by decreasing concentrations of alcohol. Sections were rinsed in tap water and then incubated in haematoxylin for min. Sections were washed in running tap water, dipped in acid alcohol (0.5 % HCl in 70 % ethanol) and then incubated in eosin for 5 min. Sections were washed in running tap water and then dehydrated in increasing concentrations of alcohol. Finally, sections were incubated in xylene for 5 min and coverslips applied using Pertex mounting medium (Cell Path Ltd, UK). Images of stained sections were captured using a Nikon N1 upright microscope (Nikon, Surrey, UK)

### Flow cytometry

Dissected brain samples were finely minced by scalpel blade in ice-cold Hanks Balanced Salt Solution HBSS (Sigma-Aldrich), centrifuged (400 g, 5 min, 4°C) resuspended and incubated for 1 h at 37°C in digestion cocktail (50 U/ml collagenase, 8.5 U/ml dispase, 100 ug/ml Nα-Tosyl-L-lysine chloromethyl ketone hydrochloride, 5 U/ml DNaseI in HBSS). Tissue was dissociated using a Dounce homogeniser and the enzymatic reaction terminated by addition of equal volume HBSS containing 10% fetal bovine serum. Homogenates were centrifuged (400 g, 5 min, 4°C) and pellets resuspended in 35% Percoll (GE Healthcare, Sweden), overlaid with HBSS then centrifuged (800 g, 45 min, 4°C). The cell pellet was resuspended in FACS buffer (DPBS containing 0.1% low endotoxin BSA). Cell suspensions were added to a 96-well plate and low affinity Fc receptors blocked by incubation of cells with anti-CD16/32 antibody for 30 min. Plates were centrifuged (400 g, 5 min), supernatants discarded and cell pellets disrupted by gentle agitation of plates. Cells were incubated with the following fluorochrome-conjugated rat monoclonal antibodies for 30 min: antimouse F4/80 (clone BM8, 1 μg/ml, BioLegend); anti-mouse/human CD11b (clone M1/70, 1 μg/ml, BioLegend); anti-mouse CD45 (clone 30-F11, 1 μg/ml, BioLegend), anti-mouse Ly6G (clone 1A8, 1 μg/ml, BioLegend) and anti-mouse CD3 (clone 17A2, 1 μg/ml, BioLegend). Plates were centrifuged (400 g, 5 min), supernatants discarded and cells resuspended in FACS buffer. Data were acquired using a LSR Fortessa (BD Biosciences) and analysed using Summit (Dako) or FlowJo software. Compensation was performed using single-labelled samples as references and positive regions of staining were defined based on unstained. Gating and criteria for identifying discrete cell subpopulations were as described in the Results sections. The absolute number of cells of a specific subset was determined from the number of cells in the population divided by total viable cells, multiplied by the total viable cell yield from the hemisphere (derived from cell density calculated by haemocytometer on the single cell suspension). Mean fluorescent intensities (MFI) of selected cell populations were derived directly from software.

### RNA extraction for microarray

RNA was extracted from a 4 mm brain sample centred on the injection site dissected using a mouse brain matrix from the hemisphere ipsilateral to injection at indicated timepoints using the miRNeasy Mini kit (Qiagen) according to manufacturer’s instructions. RNA quantities were determined by Nanodrop 1000 (Thermo Fisher Scientific, MA, USA) and RNA quality assessed using the Agilent Bioanalyzer (Agilent Technologies, CA, USA) and all samples passed a quality control threshold (RIN ≥ 7) to proceed to gene expression profiling. For microarrays, each biological replicate (n = 3) contained tissue pooled from three mice in a randomised manner and containing equivalent quantities of RNA from each constituent sample

### RNA extraction for qPCR

Unpooled samples were used for qPCR assays. Isolated mixed cell suspensions prepared as above for flow cytometry were stored in RNAlater at −80 °C. RNAlater was removed and cells were resuspended in 0.75 ml TRIzol, mixed to lyse cells, and incubated at room temperature for 5 min. 0.2 ml 1-Bromo-3-chloropropane was added to samples and shaken vigorously for 15 s, then incubated for 3 min at room temperature. Samples were centrifuged at 12,000 × *g* for 15 min at 4 °C, and upper aqueous phase was collected. Another 0.2 ml 1-Bromo-3-chloropropane was added and repeated. 1 μl of neat linear acrylamide and 0.5 ml 100 % isopropanol were added to the aqueous phase and incubated at room temperature for 10 min. Samples were centrifuged at 12,000 × *g* for 10 min at 4 °C. Supernatant was removed and the pellet was washed in 1 ml 75 % ethanol. Samples were centrifuged at 7,500 × *g* for 5 min at 4 °C, and pellet was then air-dried at room temperature. The pellet was resuspended in 30 μl RNase free water, incubated at 60 °C for 10 min and quantity determined by Nanodrop 1000.

### Transcriptomic profiling and computational analysis

Microarray assays were performed by Edinburgh Genomics, University of Edinburgh (https://genomics.ed.ac.uk/) as described previously^47^. Total RNA was labelled using the IVT Express Kit (Affymetrix). First-strand cDNA was synthesised and converted to double-stranded DNA template for transcription and synthesis of aRNA incorporating a biotin-conjugated nucleotide. aRNA was purified and fragmented prior to hybridisation on Affymetrix arrays. Biotin-labelled aRNA was hybridized to the whole mouse genome HT MG-430 PM array plate (Affymetrix, CA, USA) representing >39,000 transcripts, using the GeneTitan multi-channel instrument (Affymetrix). Microarray datasets were normalised by the Robust Multiarray Averaging (RMA) method in Affymetrix Expression Console (Affymetrix, CA, USA) and analysis performed using the ht_mg-430_pm.na36 annotation release (Affymetrix). Genes differentially expressed among treatment groups were determined by ANOVA with FDR correction using Affymetrix Transcriptome Analysis Console (TAC) and differentially expressed genes visualised by heat map. Transcripts significantly altered in expression (FDR *q* < 0.05, fold change ≥ 1.5) in LPS compared to PBS-treated mice were determined in TAC and log-transformed data visualised by volcano plot. For gene coexpression network analysis, a pairwise transcript-to-transcript coexpression matrix was computed in Graphia Pro (https://kajeka.com/graphia-professional/; Kajeka, UK) from the set of differentially-expressed transcripts among all treatments (determine in TAC as above) using a Pearson correlation threshold *r* ≥ 0.95. A network graph was generated where nodes represent individual probesets (transcripts/genes), and edges between them the correlation in expression pattern across treatment groups. The graph was clustered into discrete groups of transcripts sharing similar expression profiles using the Markov clustering algorithm in Graphia inflation = 2.2). Transcript content of key clusters was inspected manually. Cluster enrichment for Gene Ontology (GO) biological processes and cell components was determined in DAVID (https://david.ncifcrf.gov/) using default criteria.

### Quantitative PCR

cDNA was synthesised from DNase-treated RNA samples using Superscript III Reverse Transcriptase (Life Technologies) according to manufacturer’s instructions. Primers were designed using Primer BLAST (https://david.ncifcrf.gov/) and validated to ensure adequate efficiencies. Primer sequences were as follows: *Tremi1* forward 5’-CTGGTGGTGACCAAGGGTTC-3 ‘; *Trem1* reverse 5’-CTTGGGTAGGGATCGGGTTG-3’; *Trem2* forward 5’-CTGCTGATCACAGCCCTGTCCCAA-3’; *Trem2* reverse 5’-CCCCCAGTGCTTCAAGGCGTCATA-3’; *Gapdh* was used as the housekeeping gene. cDNA was mixed with forward and reverse primers, ROX reference dye and Platinum SYBR Green qPCR Supermix-UDG in DNase/RNase-free water and qPCR cycles performed on a Stratagene Mx3005P thermocycler (Agilent) as follows: hot-start denaturation cycle 95 °C for 10 min, 40 cycles of amplification at 95 °C for 15 s, 60 °C for 20 s and 72 °C for 1 min, followed by one cycle of 95 °C for 1 min, 55 °C for 30 s and 95 °C for 30 s. Cycle threshold (Ct) values of target genes were normalized to *Gapdh* and data are expressed as fold change relative to control group (PBS) using the 2^(^−ΔΔ^Ct) method.

### Experimental design and statistical analysis

Estimates of sample size were determined from previous data^31^ to achieve a power of 0.8 at P < 0.05. Data were analysed using GraphPad Prism v6 (GraphPad Software Inc, CA, USA). One-way ANOVA with post-hoc correction was used for comparisons across multiple groups (Bonferroni) or to compare selected groups to control (Dunnett). For quantitative PCR, data were analysed using a one sample *t* test compared to PBS (reference value of 1 for the PBS group). Analysis was performed unware of allocation to experimental group however no formal blinding procedure was used. Data are presented as mean ± SEM unless otherwise stated. * denotes P < 0.05, ** denotes P< 0.01, *** denotes P < 0.001, and **** denotes P < 0.0001. *P* ≤ 0.05 was considered statistically.

## Results

### Acute intracerebral LPS challenge induces a self-limiting inflammatory response with demarcated induction and resolution phases

Neutrophils are defining cells of innate immune reactions and in tissues outside the CNS, the kinetics of their accumulation and loss in response to inflammatory stimuli, such as LPS, has been useful to operatively define different phases of the inflammatory response^48^. We first assessed the effects of escalating doses of intrastriatal LPS injection on neutrophil accumulation 24 h after injection by immunohistochemistry (**Figure 1A-C**). As expected, in vehicle-injected mice there were negligible neutrophils evident in brain tissue. All doses of LPS induced marked accumulation of neutrophils in the brain parenchyma as well as increased numbers of neutrophils localised to the luminal and abluminal surfaces of blood vessels. Neutrophils were distributed throughout the striatum and the overlying motor and somatosensory cerebral cortex (**Figure 1B**). There was a concentration-dependent increase in the total number of neutrophils accumulating in brain tissue incorporating parenchymal and vascular-associated cells) 24 h after LPS injection (**Figure 1C**). For further studies, we selected the highest concentration of LPS (5 μg/μl) given the robust and consistent response. We next assessed the temporal profile of neutrophil recruitment after LPS injection. As above, LPS induced marked neutrophil accumulation 24 h after injection and similar numbers were evident at 72 h (**Figure 1D, E**). In contrast, by 5 d after injection there were few neutrophils remaining and a negligible number of cells was also evident at 7 d suggesting the infiltration had almost completely resolved by this time-point. To corroborate and extend this profile we used flow cytometric quantification of brain cell suspensions. Neutrophils were clearly identified as a distinct population of CD11b^+^Ly6G^+^ cells (**Figure 1F**). Consistent with immunostaining data, there were few neutrophils in vehicle-injected mice and a marked increase in numbers was detectable at 24 h after injection that peaked at 72 h (**Figure 1G**). Thereafter, neutrophil numbers declined markedly by 5 d and further toward control levels by 7 d where they remained stable until 28 d after injection. Using the above data we could operatively define different phases of the evolving inflammatory reaction, including a well demarcated resolution phase (**Figure 1H**). We defined the induction/amplification phase as the initial 24 h period after injection, the plateau phase from 24 - 72 h reflecting the largely stable levels of neutrophils, and the resolution phase from 72 h onwards which is reflective of the decline in neutrophils from peak levels towards baseline. Thus, intrastriatal injection of LPS induces an acute self-limiting inflammatory reaction with well-demarcated induction and resolution phases as defined according to intracerebral neutrophil accumulation and loss.

### Temporal changes to the cerebrovasculature and microglia/macrophages in relation to induction and resolution phases

Changes to the adhesive properties and permeability of the vascular endothelium as well as reactivity in tissue macrophages are archetypal features of inflammatory responses. We performed an initial qualitative assessment of selected markers to understand how gross changes to the cerebrovasculature and microglial/macrophage population evolved in relation to the phases of inflammation defined above. Immunoreactivity of ICAM-1 (**Figure 2A**), an adhesion molecule induced in inflamed endothelium that mediates leukocyte transmigration, was undetectable in vehicle-injected brain but was induced 24 h after LPS injection. Further induction, reflected by greater intensity of immunostaining, occurred at 5 d and thereafter there was declining expression. Permeability of the blood-brain barrier (BBB), a key property of the brain microvascular endothelium, was assessed by leakage of endogenous IgG, which is normally prevented from entering the brain by the BBB. Alterations in IgG immunostaining were largely confined to the injected hemisphere (**Figure 2B**). A focal area of IgG immunoreactivity within the striatum was evident 24 h after LPS injection and this extended to the entire striatum and most cortical areas at the coronal plane of injection 3 d - 7 d after injection. IBA1 immunostaining was used to assess the general pattern of microglia/macrophage reactivity and in vehicle-injected brains immunoreactivity was characteristic of ramified microglia evenly distributed throughout the parenchyma (**Figure 2C**). The intensity of IBA1+ cells within the ipsilateral striatum and cortex increased 24 h after LPS injection and there was evidence of hypertrophy of cell bodies indicative of microglial activation. At later time-points (3 −7 d), there were modest increases in the density of IBA1+ cells (see flow cytometry data below for quantification) and a markedly greater range of cell morphologies and intensities evident. This included ramified cells with an intense, hypertrophic cell soma similar to those observed at 1 d and the appearance of cells with thickened, short processes and much larger, very intensely-stained cells largely devoid of processes (**Figure 2C**). This pattern indicated an increasing heterogeneity of the IBA1^+^ population from 3 d onwards coinciding with the resolution phase and likely reflected reactive changes in resident microglia and the accumulation of monocyte-derived macrophages. We also noted that there was negligible if any signs of microglial reactivity in vehicle-injected mice suggesting the mechanical nature of the injections themselves was not an inflammatory trigger and consistent with the absence of histological signs of trauma (**Figure 1D**).

**Figure 2.**
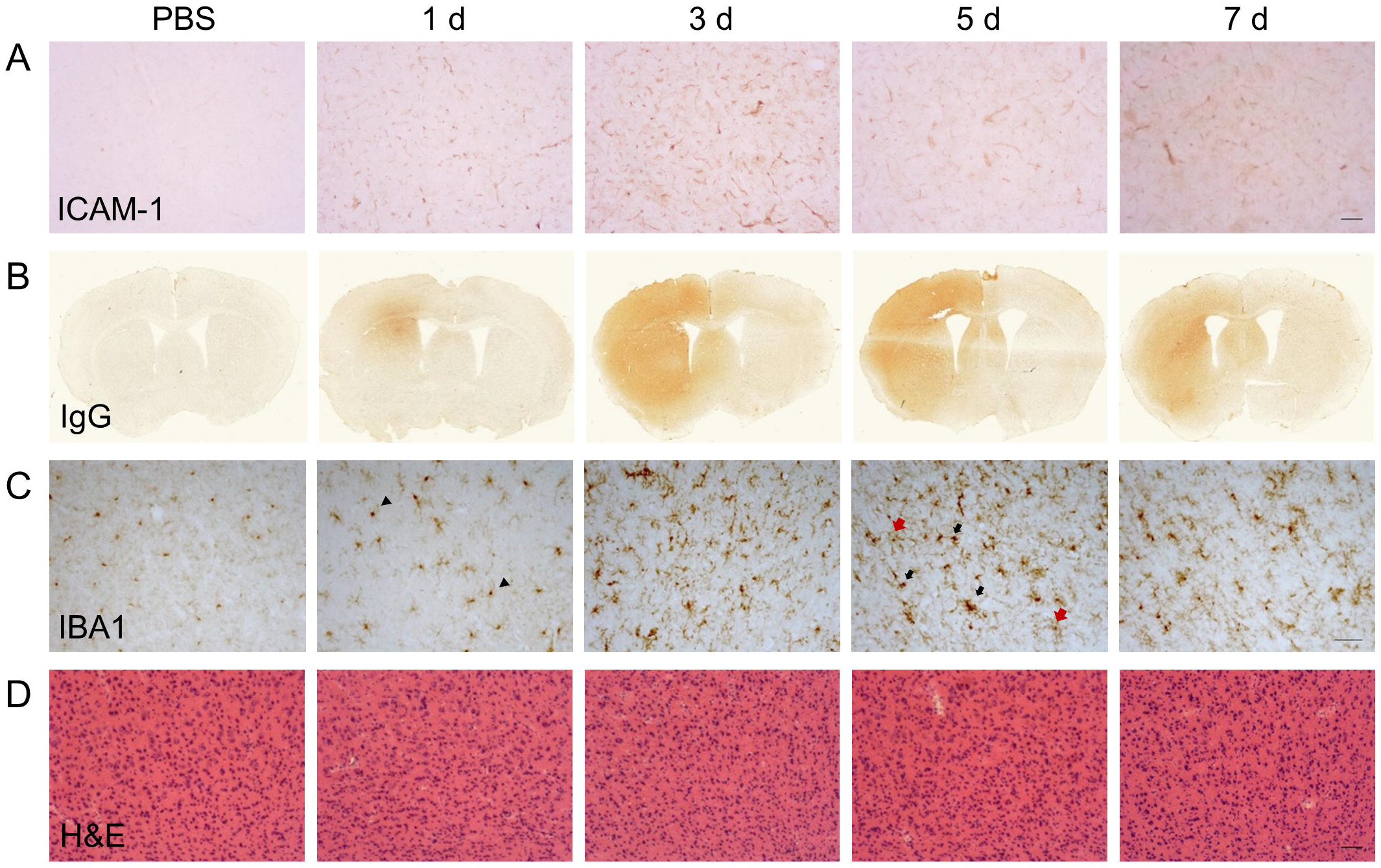
Temporal profile of cerebrovascular and gross CNS macrophage changes after intracerebral LPS challenge. Representative images of (A) cortical ICAM-1 and (B) IgG immunostaining at indicated timepoints after intracerebral LPS challenge indicate changes in cerebrovascular activation and blood-brain permeability. (C) Representative images of cortical IBA1 immunostaining at indicated timepoints after intracerebral LPS challenge. Morphological changes in IBA1^+^ cells from ramified in PBS-treated mice to intensely IBA1^+^ with shorter processes (1 d, arrowheads) and hypertrophic (5 d, black and red arrows) after LPS challenge were evident. (D) Representative images of haematoxylin and eosin (H & E) staining indicated no overt neuronal pathology caused by LPS injection. Scale bar: 100 μm (A, D); 50 μm (C).

### Accumulation of monocyte-derived macrophages coincides with the onset of inflammation resolution

In view of the immunohistochemistry observations suggestive of an evolving mixed microglial/macrophage population in LPS-injected brains, we used flow cytometry to investigate the myeloid cell composition more thoroughly and quantitatively. Combinatorial expression of a panel of myeloid cell markers (CD11b, Ly6G, CD45) enabled identification of distinct subsets. In vehicle-injected brains a relatively homogenous population of CD11b^+^CD45^lo^Ly6G^−^ microglia was evident whereas after LPS injection additional CD11b+ populations appeared (**Figure 3A**). These included Ly6G+ neutrophils (see above) and two major subsets of CD11b^+^Ly6G^−^ cells that could be distinguished on the basis of differential CD45 expression intensity. We operationally defined CD11b^+^Ly6G^−^CD45^lo^ cells as microglia i.e. resident macrophages of the brain parenchyma) and CD11b^+^Ly6G^−^CD45^hi^ cells as recruited monocytes (Mo)/monocyte-derived macrophages (Mo/MDMs) (a small proportion may comprise perivascular and/or choroid plexus macrophages) (**Figure 3A**). Although microglial CD45 expression can be induced in the inflamed state, previous studies have shown that differential CD45 intensity reliably distinguishes these different sources. Indeed, in the present study at all time-points after LPS injection, mean CD45 intensity on the CD11bLy6G^−^CD45^hi^ population was 5-10-fold greater including when microglial CD45 intensity was induced to its maximum (almost doubling) at 3 d (**Figure 3B**, 4B). The number of total CD11b+ cells increased in LPS-injected brains reaching a peak 3 d after injection and declining to near baseline levels by 28 d (**Figure 3C**). The number of CD11b^+^Ly6G^−^CD45^lo^ microglia did not significantly increase in response to LPS challenge (**Figure 3D**). Negligible numbers of CD11bLy6G^−^CD45^hi^ cells were present in vehicle-injected brains and a marked increase in number was evident after LPS challenge with maximal accumulation 3 d after injection and a gradual decline thereafter (**Figure 3E**). To corroborate these findings and ensure there was no misidentification of microglia and Mo/MDMs using differential CD45 expression, we performed a separate experiment using heterozygous *Ccr2*^RFP+^ reporter mice that express RFP in monocyte-derived cells but not microglia. RFP^+^ cells were largely absent from vehicle-injected brains whereas a clear accumulation of RFP^+^ cells was observed 3 d after LPS injection (**Figure 3F**). RFP expression was only found in Ly6G^−^ cells that were also expressing high levels of CD45 and not in CD11b^+^CD45^lo^ cells and almost all CD11b^+^CD45^hi^ cells were positive for RFP expression (**Figure 3G**). These data confirm the influx of Mo/MDMs and validate the use of differential CD45 expression to distinguish resident and recruited CNS macrophages in this model. This also excludes the theoretical possibility that the increase in number of CD11bLy6G^−^CD45^hi^ cells in response to LPS is predominantly an expansion of the perivascular macrophage population (which express higher levels of CD45 compared to parenchymal microglia) because previous studies have shown these are not derived from and do not turnover from CCR2-dependent (RFP) bone marrow precursors^49^. Overall, the flow cytometry data are consistent with the immunohistochemistry observations of a heterogeneous microglial/macrophage population and show that a marked accumulation of recruited Mo/MDMs coincides with the onset of resolution 3 d after LPS injection.

**Figure 3.**
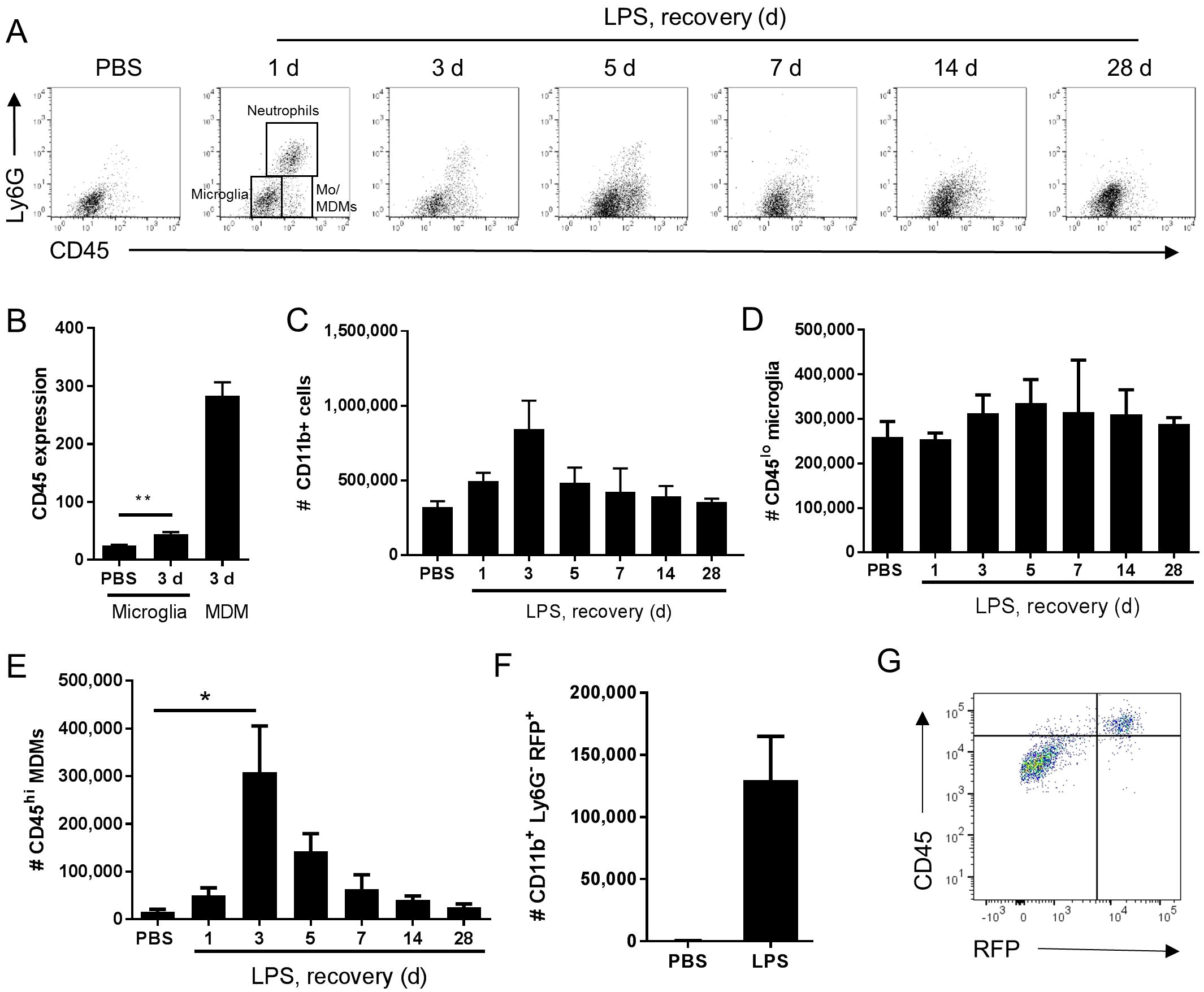
Temporal profile of microglial and monocyte-derived macrophage recruitment after intracerebral LPS challenge. Brain cell suspensions were prepared from the ipsilateral hemisphere after intrastriatal PBS or LPS injection at indicated timepoints. (A) Representative flow cytometry dot plots and regions defining selected myeloid cell subsets (annotated at 1 d timepoint after LPS) based on Ly6G^+^ and CD45^+^ labelling (with pre-gating on CD11bCD3^−^ cells). Neutrophils were defined as Ly6GCD45^hi^, microglia as Ly6G^−^CD45^lo^, and monocyte-derived macrophages (Mo/MDM) as Ly6G^−^CD45^hi^. Dot plots show representative changes in the composition of cell subsets as time progresses after LPS challenge. (B) Quantification of CD45 expression intensity in microglia 3 d after LPS with MDM expression shown for reference. **P < 0.01, Student’s *t* test. Quantification of the total number of (C) CD11b^+^ cells, (D) CD11b^+^CD3^−^ Ly6G^−^CD45^lo^ microglia, and (E) CD11b^+^CD3^−^Ly6G^−^CD45^hi^ Mo/MDMs in the ipsilateral hemisphere after PBS or LPS injection. *P < 0.05, one-way ANOVA with Bonferroni’s post-hoc test. (F) Quantification of the total number of CD11b^+^Ly6G^−^RFP^+^ MDMs after 3 d after intrastriatal PBS or LPS injection in *Ccr2*^RFP/+^ reporter mice. (G) Representative flow cytometry dot plot showing relationship between RFP and CD45 expression in brain cell suspensions from *Ccr2*^RFP/+^ reporter mice 3 d after intrastriatal LPS injection.

### The peak of microglial reactivity corresponds with the inflammation resolution phase

As described above, morphological signs of microglial reactivity were evident during both the induction and resolution phases, although appeared most prominent during resolution. Using flow cytometry, we analysed expression intensities of general markers of microglial/macrophage reactivity and maturation, CD45 and F4/80, specifically on the microglial population (CD11b^+^Ly6G^−^CD45^to^ cells as above). Mean fluorescence intensity of both microglial CD45 and F4/80 increased significantly in response to LPS injection (**Figure 4A-D**). This induction was relatively modest at 24 h after injection induction phase) but was maximal at 3-5 d thus coinciding with the resolution phase and suggesting the magnitude of gross microglial reactivity peaks during resolution (**Figure 4A-C**). Comparison of peak CD45 (3 d) (**Figure 3B**, **4B**) and F4/80 (5 d) (**Figure 4C-D**) expression levels in microglia with those in Mo/MDMs at the corresponding time-point after LPS challenge confirmed that, despite 2-fold induction of CD45 and 5-fold induction of F4/80 compared to vehicle-injected mice in microglia, expression remained markedly lower than in Mo/MDMs (~10-fold lower for CD45; ~2-fold lower for F4/80). In agreement with the above flow cytometry data, mining of microarray data (see below for details) showed that expression levels of the genes encoding CD45 and F4/80, *Ptprc* and *Emr1*, were significantly elevated in LPS-injected brain homogenates and greatest during the resolution phase (**Figure 4E**) (NB: we are not excluding that this will also partly reflect the greater number of CD45^hi^ cells that accumulate during this phase as shown in **Figure 3**). Overall, these data suggest that the resolution phase is associated with a particularly reactive state of resident microglia that occurs alongside the recruitment of Mo/MDMs.

**Figure 4.**
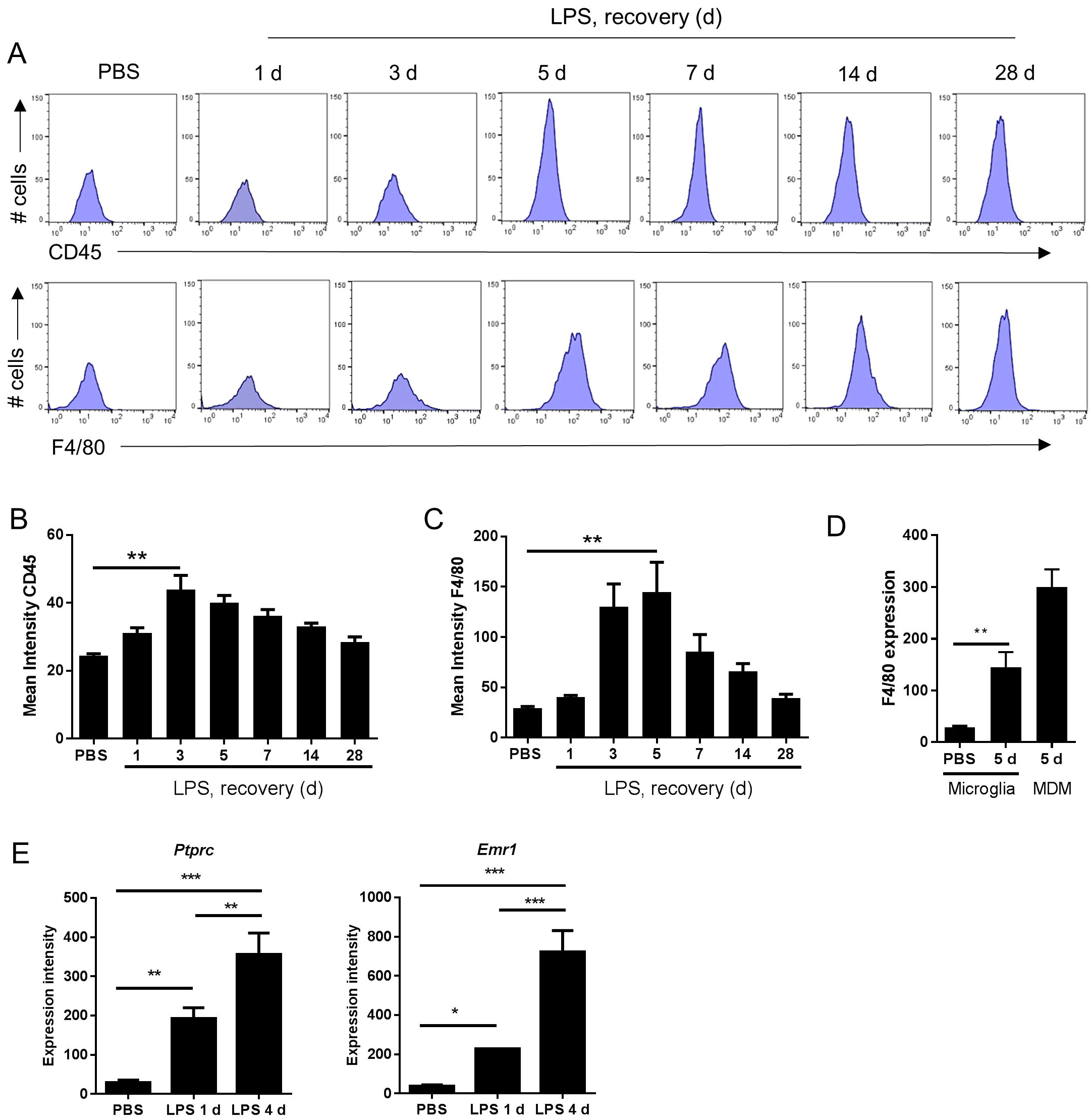
Markers indicative of the magnitude of microglial reactivity peak at the onset of and during the resolution phase after intracerebral LPS challenge. (A) Brain cell suspensions were prepared from the ipsilateral hemisphere after intrastriatal PBS or LPS injection at indicated timepoints. Representative histograms showing expression of CD45 (top panel) and F4/80 (bottom panel) in CD11b^+^Ly6G^−^CD45^lo^ microglia. Quantification of the mean fluorescence intensity of (B) CD45 and (C) F4/80 in CD11b^+^Ly6G^−^CD45^lo^ microglia. **P < 0.01, Student’s t-test (B); **P < 0.01, Mann-Whitney test (C), respectively. (D) Change in mean fluorescence intensity in CD11b^+^Ly6G^−^CD45^lo^ microglia of F4/80 5 d after LPS compared to PBS shown with CD11bLy6G^−^CD45^hi^ MDM CD45 intensity shown for reference. **P < 0.01, Mann-Whitney test. (E) Gene expression intensity of *Ptprc* and *Emr1* measured by microarray from brain homogenates isolated from the ipsilateral hemisphere after intrastriatal PBS or LPS injection at indicated timepoints. *P < 0.05, **P < 0.01, ***P < 0.001; one-way ANOVA with Bonferroni’s post-hoc test.

### Inflammation resolution is associated with *de novo* induction of a unique transcriptional programme

The above data show some of the key cellular changes associated with resolution of acute brain inflammation. We next sought to determine if inflammation resolution was associated with expression of distinct transcriptional networks by using a transcriptome-wide approach to profile gene expression at resolution and induction phases and in the uninflamed state (PBS injection). On the basis of cellular data described above, we selected 4 d after LPS (resolution) and 1 d after LPS induction) as representative of the different phases of the inflammatory response. 1779 transcripts were differentially expressed across all treatment groups (ANOVA with FDR *q* < 0.05) and a heat-map of these transcripts demonstrated distinct patterns of altered gene expression from the uninflamed state and between resolution and induction phases (**Figure 5A**). Expression of a negligible number of transcripts was suppressed 1 d or 4 d after LPS compared to PBS injection therefore we focussed on upregulated transcripts. Expression of 723 transcripts was significantly increased (FDR *q* < 0.05, fold-change ≥ 1.5) 1 d after LPS and 534 transcripts 4 d after LPS compared to PBS injection (**Figure 5B**). We identified 396 induction phase-specific and 207 resolution phase-specific transcripts respectively based on these filtering criteria (**Figure 5C**). This provided an initial indication that resolution was associated with induction of a unique *de novo* transcriptional programme.

**Figure 5.**
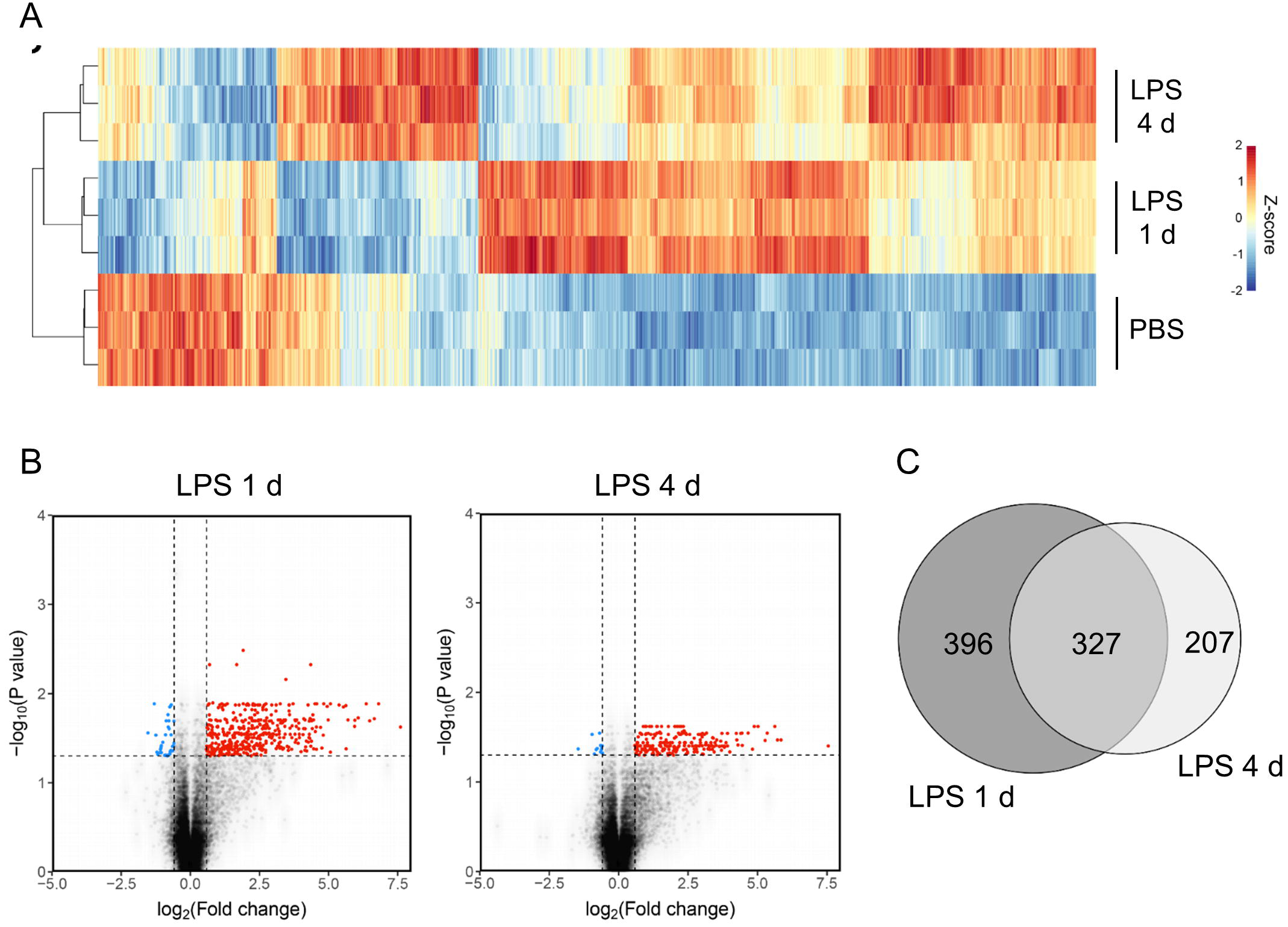
Transcriptome profiling identifies unique sets of genes induced at the induction and resolution phases of the inflammatory response after intracerebral LPS challenge. Brain homogenates were prepared from the ipsilateral hemisphere after intrastriatal PBS or LPS injection at indicated timepoints and transcriptome profiling by microarray performed. (A) Visualisation by heat map of the 1779 transcripts differentially expressed across all treatment groups (ANOVA with FDR *q* < 0.05) and hierarchical clustering of samples according to the transcript expression patterns. Note the distinct modules of transcripts selectively altered 1 d μnduction phase) and 4 d (resolution phase) after LPS injection. (B) Visualisation of the transcripts differentially expressed 1 d or 4 d after LPS compared to PBS injection (FDR *q* < 0.05, fold-change ≥ 1.5) by volcano plot. (C) Venn diagram depicting the overlapping and unique transcripts significantly induced 1 d or 4 d after LPS compared to PBS injection (FDR *q* < 0.05, fold-change ≥ 1.5).

### Co-expression network analysis and represented functional pathways of inflammation induction and resolution

To explore the composition of the transcriptional networks underpinning resolution and their differences with the induction phase in more depth we constructed a gene co-expression network graph and non-subjectively clustered into groups of highly co-expressed genes. Four major clusters of co-expressed genes dominated the network graph (clusters 1-4) (**Figure 6A**) and we therefore focussed further attention on these. The mean expression profile of each of the major clusters showed a distinct profile (**Figure 6B**): cluster 1 showed increased expression relatively selectively 1 d after LPS challenge (hereafter referred to as the induction-selective cluster); cluster 2 showed the opposite pattern with increased expression relatively selectively 4 d after LPS challenge (hereafter referred to as the resolution-selective cluster); cluster 3 showed increased expression at both 1 d and 4 d after LPS challenge with relatively greater expression at 4 d (hereafter referred to as the resolution-dominant cluster); cluster 4 also showed increased expression at both 1 d and 4 d after LPS challenge but with relatively greater expression at 1 d (hereafter referred to as the induction-dominant cluster). We manually inspected the contents of each cluster and an overall summary is presented in **Supplementary Table 1** - here we largely focus on the induction- and resolution-selective clusters summarised in **Figure 6C**.

**Figure 6.**
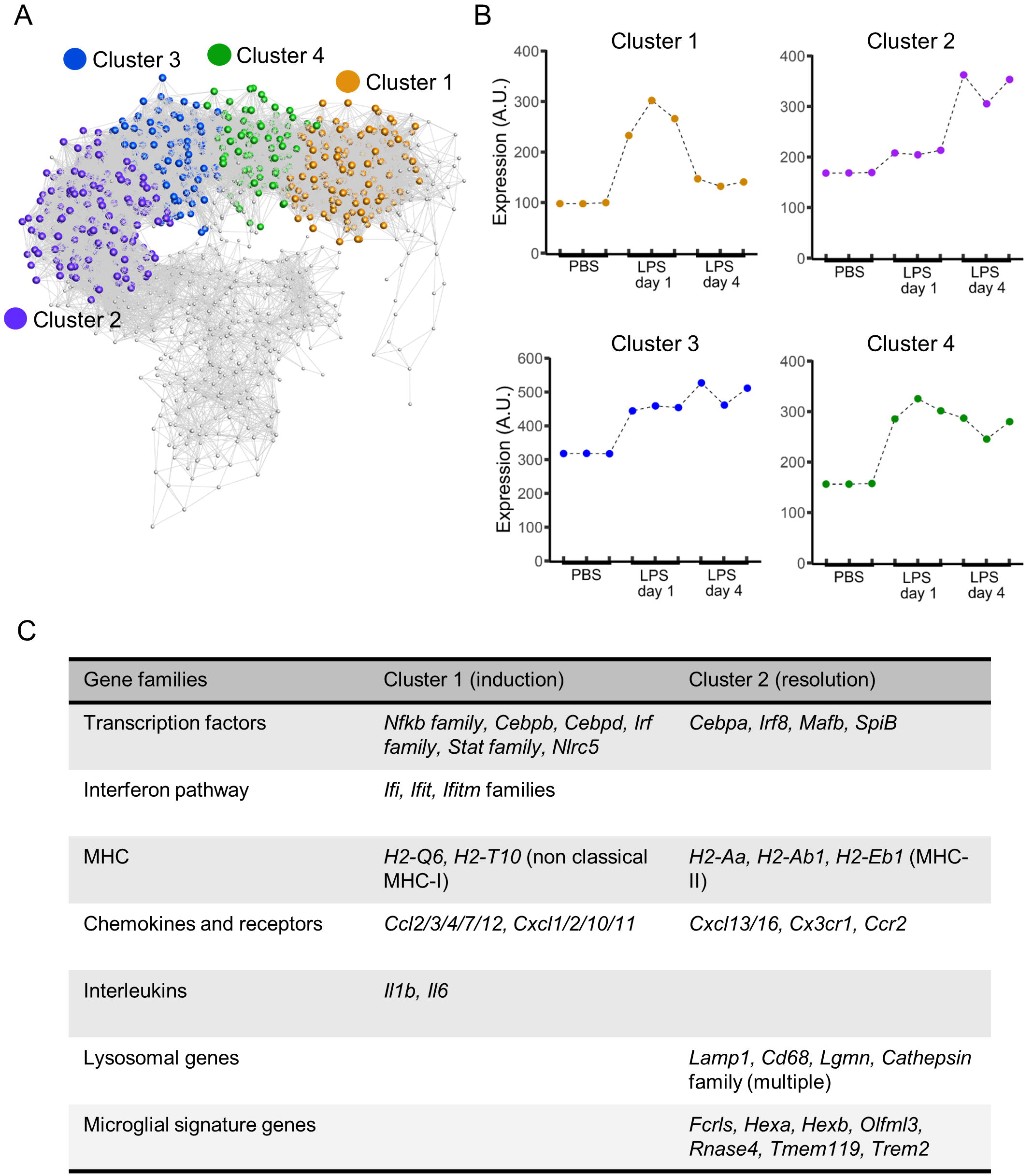
Gene coexpression analysis of transcriptome data after intracerebral LPS challenge shows distinct clusters of coexpressed genes including a resolution-specific cluster. Brain homogenates were prepared from the ipsilateral hemisphere after intrastriatal PBS or LPS injection at indicated timepoints and transcriptome profiling by microarray performed. (A) Gene coexpression network graph of the 1779 transcripts differentially expressed across all treatment groups. The coexpression matrix was clustered using a Markov clustering algorithm and the major clusters of coexpressed transcripts are highlighted. (B) Mean expression profiles of clusters 1-4 across all treatment groups derived from all transcripts in each cluster. (C) Clusters were manually inspected for transcript composition. Selected gene families of interest over-represented in cluster 1 μnduction-selective) and cluster 2 (resolution-selective). For full summary of clusters 1-4 see **Supplementary Table 1**.

The induction-selective cluster (cluster 1) contained 419 nodes/transcripts. Manual inspection of the cluster revealed a number of functional classes of genes consistent with a co-ordinated role in sensing, initiation and amplification of an innate immune response (**Figure 6C**, **Supplementary Table 1**). These included Toll-like receptors (*Tlr3*), scavenger receptors (*Marco*, *Msr1*), cytokines (*Il1*, *Il6*), endothelial adhesion molecules (*Icam1*, *Vcam1*, *Madcam1*) and chemokines of both CC (*Ccl2/3/4/7/12*) and CXC classes (*Cxcl1/2/10/11*). Further chemokines (*Ccl5/6/9/19*) were present in the induction-dominant cluster (cluster 4). There was a particularly notable presence of a large number of interferon-regulated genes of the *Ifi*, *Ifit* and *Ifitm* families in the induction-selective cluster suggestive of potent interferon signalling during this phase. This was further evident by the large number of classical (*H2-D1*, *H2-K1*, *H2-Q7*) and non-classical MHC class I (*H2-Q6*, *H2-T10*) genes collectively contained with the induction-selective and -dominant clusters. In contrast, MHC class II genes were not present within the induction clusters. Consistent with this pattern of interferon and MHC class I gene expression during the induction phase, several transcription factors regulating these pathways were evident in the induction-selective cluster including from the *Irf* family (*Irf1/7/9*), the *Stat* family (*Stat1/2/3*) and the master regulator of MHC class I expression, *Nlrc5*. Other transcription factors contained within the induction-selective and -dominant clusters included genes encoding several components of NF-κB (*Rela*, *Relb*, *Nfkb2*) and from the *Cebp* family (*Cebpb*, *Cebpd*) that are consistent with induction of a range of inflammatory mediators described above. In support of the above observations, GO enrichment analysis revealed over-representation of the biological processes “chemotaxis”, “response to interferon-alpha”, “response to interferon-beta “, and “antigen processing and presentation of exogenous peptide antigen via MHC class I” (**Supplementary table 2**). Collectively, these data indicate a potent innate immune transcriptional programme during the induction phase featuring a prominent interferon and MHC class I response among an array of broader inflammatory signalling.

The resolution-selective cluster (cluster 2) contained 347 nodes/transcripts and comprised groups of functionally-related genes distinct from those in the induction clusters emphasising the engagement of transcriptional networks specific to the resolving phase (**Figure 6C**, **Supplementary Table 1**). In contrast to the induction clusters, which contained a prominent MHC class I gene set (see above), genes encoding proteins involved in antigen processing and presentation on MHC class II (*H2-Aa*, *H2-Ab1*, *H2-Eb1*, *Ifi30*, *Cd74*) were selectively present in the resolution-selective cluster suggestive of differing involvement of the type of antigen processing and presentation as the inflammatory response evolves. Of particular note, an array of genes encoding lysosomal proteins was evident in the resolution-selective cluster which included the lysosomal membrane protein-encoding *Lamp1*, *Laptm5* and *Cd68* and a range of protease-encoding genes such as the cathepsins (*Ctsa/b/h/s/z*), glycosidases (*Gusb*), hexosaminidases (*Hexa*, *Hexb*), legumain (*Lgmn*), lipases (*Lipa*) lysozymes (*Lyz1*, *Lyz2*) and sulfatases (Gns). This set of genes is indicative of enhanced lysosomal activity and phagocytosis and further highlights the induction of MHC class II pathway given the important role for endo-lysosomal cathepsins in peptide processing and loading for antigen presentation^50^. Further functional classes of genes of interest in the resolution-selective cluster included lipid metabolism (*Apobec1*, *Apoc2*, *Apoe*, *Lipa*, *Npc2*) and the complement pathway (*C1qa/b/c*, *C3ar1*). In support of the above findings, GO enrichment analysis revealed over-representation of the biological processes “antigen processing and presentation of exogenous peptide antigen via MHC class II”, “lipoprotein biosynthetic process”, “cholesterol efflux”, “proteolysis involved in cellular protein catabolic process”, and “phagocytosis, engulfment” (**Supplementary table 3**) among general immune response enrichment as well as the cellular component “lysosome” (**Supplementary table 4**). It was also evident that a distinct set of transcription factors, including *Cebpa*, *Irf8*, *Mafb* and *Spib* were present in the resolution-selective cluster which is consistent with regulation of myeloid cell differentiation/maturation and the flow cytometry data described above. Overall, these data reinforce the distinctive nature of the regulatory and transcriptional networks controlling the resolution phase and in particular suggest lysosomal function as central to multiple co-ordinated pro-resolution molecular processes.

### Inflammation resolution involves induction or restoration of homeostatic microglial gene expression

Recent studies examining the transcriptional basis of microglial identity have described a set of genes highly enriched in microglia compared to systemic myeloid cells and other CNS cell types^47,51–54^. This expression signature has been proposed to reflect a homeostatic microglial state^55^. We noticed that the resolution-selective cluster contained a considerable number of these microglial signature genes, including *Fcrls*, *Hexa*, *Hexb*, *Olfml3*, *Rnase4*, *Tmem119*, and *Trem2* (**Figure 6C**). The expression of all these genes was significantly increased 4 d after LPS challenge compared to 1 d and also after PBS treatment (**Figure 7A**). Expression of *P2ry12* and *P2ry13*, further proposed homeostatic microglial signature genes, was also elevated 4 d compared to 1 d after LPS challenge (**Figure 7B**) although this largely reflected a restoration of suppressed expression levels and is consistent with the known instability of these genes in response to LPS or other overtly pro-inflammatory conditions. We did not observe changes in microglial numbers during the transition from induction to resolution phases (see **Figure 3** above) therefore gene expression changes are likely the result of actual quantitative expression changes. *Trem2* is a member of a receptor family that in mice also contains *Trem1* and *Trem3.* In contrast to *Trem2,* expression of *Trem1* and *Trem3* was selectively induced 1 d after LPS challenge (**Figure 7C**) in keeping with their presence in the induction-selective cluster and known functionally distinct role from *Trem2* in amplification of innate immune signalling^56^. We validated the microarray expression pattern of the *Trem* genes by qPCR on myeloid cell-enriched cell suspensions isolated from the brain over an extended time-course after LPS challenge. This showed the same opposing pattern of *Trem1* and *Trem2* expression and with similar kinetic profile as microarray analysis (**Figure 7D**). Overall, these data suggest that resolution of acute brain inflammation is associated with induction and/or restoration of “homeostatic” microglial signature genes and a transition in the balance of amplifying and modulatory immune signalling via immunoreceptors such as TREM1 and TREM2.

**Figure 7.**
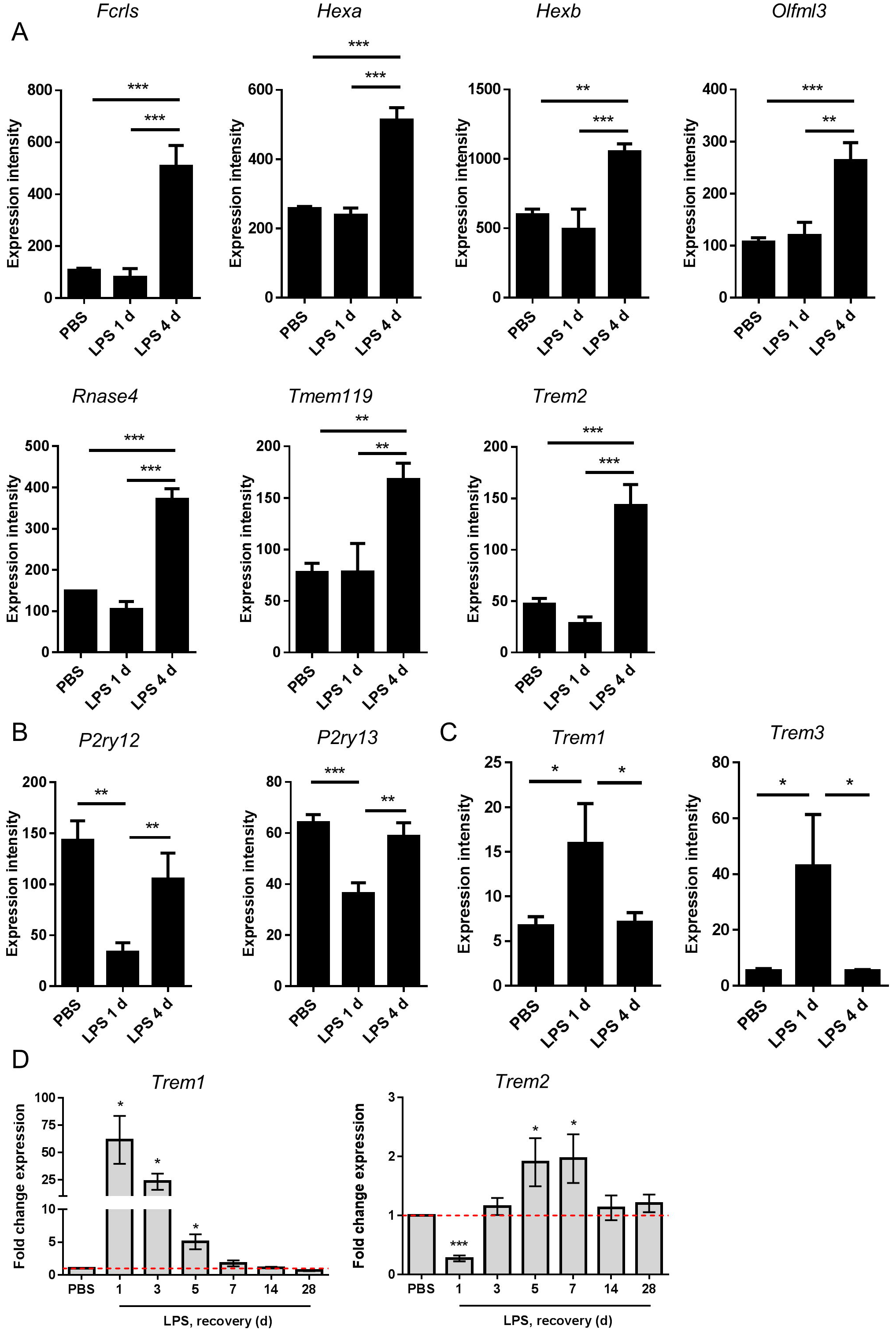
Changes in expression of microglial signature genes after intracerebral LPS challenge suggest restoration of a homeostatic microglial state is associated with inflammation resolution. Brain homogenates were prepared from the ipsilateral hemisphere after intrastriatal PBS or LPS injection at indicated timepoints and gene expression measured by (A, B) microarray or (C) quantitative PCR. Expression intensities of (A) selected microglial homeostatic signature genes, (B) P2Y receptor genes, and (C) amplifying TREM family genes. *P < 0.05, **P < 0.01, ***P < 0.001; one-way ANOVA with Bonferroni’s post-hoc test. (D) Relative expression of *Trem1* and *Trem2* after LPS versus PBS injection at indicated timepoints. *P < 0.05, one sample *t* test compared to PBS (reference value of 1 for the PBS group).

## Discussion

In the present study, we sought to develop and validate a model tailored for studying the resolution of acute brain inflammation and in doing so define its key cellular and molecular characteristics. Several interesting observations emerged, notably that the resolution phase appears to be a very active period of transcriptomic reprogramming temporally aligned with marked changes in myeloid cell composition and reactivity. The very dynamic nature of this phase points to its potential for manipulation and subversion thus understanding endogenous regulatory mechanisms and factors perturbing them will be important in future. The baseline data provided here provide a platform for future studies to uncover novel CNS resolution mechanisms and modifiers.

We used LPS as the inflammatory challenge which has both strengths and weaknesses. As a cell wall component of gram-negative bacteria, LPS is relevant to infectious challenges of the brain. Although rare, infection of the central nervous system parenchyma often has severe consequences^57^ and it is important to consider previous studies that indicated an unrestrained host inflammatory response to infection may be more problematic than direct pathogen-induced toxicity^58^. LPS is also likely relevant to broader contexts of sterile inflammation given its actions are mediated by signalling through TLR4. Other ligands for TLR4 include a number of damage-associated molecular patterns (DAMPs), such as HMGB1 and S100A8/9, which are produced and/or exposed by cell death and other triggers of sterile inflammatory responses important in the reaction to brain hypoxia/ischaemia, trauma^59,60^. Microbial and sterile challenges can result in conserved TLR4 signalling responses (e.g. MAP kinase, NF-κB and HIF-1 activation) that induce closely overlapping transcriptional and metabolic reprogramming profiles. Indeed, our unpublished data suggest a substantial degree of overlap in the transcriptome-wide changes in gene expression in response to intracerebral LPS injection or experimental stroke in mice. In the present study there was no evidence that LPS injection caused neuronal toxicity therefore the initial inflammatory response should reflect a direct response to LPS thus exposure to a highly consistent stimulus is achieved. In most models of acute brain injury, cell death is a key feature but often is variable in magnitude and distribution. In some cases, it will be necessary to include the complexity of processes inherent in acute brain injuries (e.g. cell death, oedema, blood flow alterations) but in other instances it may be advantageous to use a more reductionist but nonetheless representative system as described here.

Mechanisms of inflammation resolution have been studied extensively in tissues outside the brain. The utility of reductionist models is emphasised by their leading to the discovery of novel pro-resolving mediators, notably several classes of fatty acid-derived molecules^61^. Models used have included the intraperitoneal injection of LPS or zymosan and the air-pouch inflammation model^48,62,63^. Key features of these models are the ability to analyse compartmentalised tissue/fluid extracts with precise monitoring of kinetic profiles of molecular and cellular changes. In common with these models, we observed a range of cellular changes characteristic of an acute inflammatory response regardless of tissue location, including resident macrophage activation, increased vascular permeability and endothelial activation, and myeloid cell recruitment consisting of early neutrophil accumulation and a slower influx of monocyte-derived cells. Detailed analysis in a murine acute peritonitis model produced quantitative indices for the temporal evolution of key cellular changes, including the resolution index based on the relationship between neutrophil and monocyte accumulation^48^. In the present study, neutrophil accumulation peaked 1-3 d after LPS injection and there were negligible numbers by 5 d indicating a relatively rapid removal of these cells from the brain. We estimated the resolution index (time for neutrophil numbers to decrease to half their peak value) to be 18 h in the present intracerebral LPS model. While this shows an ability of the brain to clear a substantial influx of neutrophils rapidly, it is interesting to consider this is likely slower than described for analogous systemic models^48^. Thus, while there are a number of shared features of acute inflammation triggered inside and outside the brain, there are also likely location-dependent properties as we recently demonstrated^31^.

An important conceptual principle to emerge that underpins how inflammation is brought under control outside the brain is that it involves active induction of pro-resolution cellular and molecular processes and is not simply the waning of the initial inflammatory response^64^. In the present study, there are several observations that support a highly active resolution phase in the acutely inflamed CNS. This includes the temporal pattern of CNS macrophage accumulation and reactivity. Using differential CD45 expression by flow cytometry we were able to identify both resident microglial and infiltrating monocyte sources of CNS macrophages in the LPS-injected brain, which importantly we validated the specificity of using *Ccr2*-RFP monocyte reporter mice. Infiltration of monocyte-derived macrophages peaked 3 d after LPS coinciding with the onset of resolution. At a similar time (3-5 d after LPS), microglial numbers peaked although we note this involved only a modest and statistically non-significant increase. During this time-frame, the most visible morphological signs of CNS macrophage IBA1^*+*^) reactivity were also observed. Furthermore, surface protein expression levels of CD45 and F4/80 specifically on microglia steadily increased to reach a peak 3-5 d after LPS again coinciding with the resolution phase and loss of neutrophils. Expression of the genes encoding CD45 (*Ptprc*) and F4/80 (*Emr1*) followed a similar pattern. CD45 and F4/80 are induced by microglial/macrophage activation and/or differentiation and across the time-course studied here provide an indication of relative temporal changes in magnitude of gross microglial activation. Together, these data indicate that accumulation and indices of reactivity of CNS macrophages reach their maximum during the onset and progression of resolution. Mononuclear phagocytes have previously been shown to participate in several aspects of a resolving inflammatory process including through efferocytosis of neutrophils, provision of trophic and growth factors and signalling to stromal and other immune cells^26,65,66^. Our data are therefore suggestive of resolution-promoting active changes in CNS macrophage composition and reactivity. Future studies targeting specific mononuclear phagocyte subsets will be required to confirm their individual and collective functional roles in resolution.

Perhaps the most striking evidence that the resolution phase in the present model involves actively induced cell and molecular programmes comes from the transcriptomic profiling identifying distinct gene signatures associated with inflammation induction and resolution. Using an unbiased gene coexpression network approach, we identified a cluster of coexpressed genes with largely selective (or substantially greater) induction in expression during the resolution phase thus confirming active *de novo* gene network reprogramming as an integral CNS resolution event. A number of molecular classes of genes were contained within this cluster and their coexpressed pattern implicates likely coordinated functions of their encoded proteins and co-regulation by common transcription factors. Here, we discuss some particularly notable features of the resolution gene cluster and their contrasts with the induction phase cluster (see **Supplementary Table 1** for overview of wider cluster composition and associated biology for all gene clusters).

One of the most prominent classes of genes in the resolution cluster was the lysosomal group that included genes encoding multiple cathepsins, lysozyme, several other lipid-targeted proteases and lysosomal membrane proteins. Lysosomal function is critical to phagocytic removal of cells and debris, and our data may reflect this role in the rapid loss of neutrophils by mononuclear phagocytes in the current model. Lysosomes also act as a convergence point for autophagy- and phagocytosis- derived intra- and extra-cellular materials, and pro-resolving functions of macrophages, including phagocytosis, were associated with enhanced autophagy^67^. A further important function for lysosomal proteases is the processing of peptide antigens for presentation on MHC class II^50^. Our data showed a marked switch in the classes of MHC genes induced from the induction (MHC-I) to resolution (MHC-II) clusters. This may be similar to the enrichment of resolution-phase macrophages in the peritoneum for MHC-II-related genes^68^ and suggests antigen presentation and links between innate and adaptive immune processes may be important for resolution^66^. Although we did not assess lymphocyte subsets in the current study, increased expression of several T cell and B cell chemoattractants in LPS-challenged mice was evident in our transcriptome dataset. Overall, the array of lysosomal proteases induced during resolution may therefore have multiple functions.

The resolution cluster also contained a large group of genes encoding proteins involved in lipid scavenging and metabolism, particularly apolipoproteins and including the major lipid transport protein of the CNS, apolipoprotein E^69^. The increased expression of these mediators may be important to provide increased capacity for sequestering, processing and recycling lipid components of dying or phagocytosed cells and it is logical to observe coexpression of these genes alongside the lysosomal genes described above. *Cd5l*, part of the resolution cluster, and a gene that lies at the intersection of lipid metabolic and immune regulation^70^ further highlights how altered lipid metabolism may be an integral component of successful inflammation resolution.

The resolution cluster contained several transcription factors associated with reactivity and differentiation of microglia/macrophages (e.g. *Ets1*, *Irf8*, *Mafb*, *Spib*)^71–73^ consistent with cellular data discussed above and suggesting possible key control hubs of resolution gene networks, including those within the resolution cluster. The identity of these resolution-phase transcription factors was particularly noteworthy when contrasted with the induction cluster. Genes encoding NF-κB, interferon response factors IRFs) and STAT proteins were prominent in the induction cluster and in keeping with the array of MHC-I, interferon and interleukin genes induced at the induction phase that are known targets of these transcriptional regulators. Thus, our data highlight not only marked shifts in multiple functionally-linked gene networks as the inflammatory response transitions to resolution but also in the upstream transcriptional control pathways that are likely governing this transition.

We and other groups have recently described transcriptome-wide profiles defining the identity and diversity of rodent microglia in the steady-state^47,52,53^. Expression of a unique “homeostatic” gene set is a defining feature of microglia in the non-pathologic CNS and altered expression of genes within this homeostatic cluster can indicate deviations towards reactive phenotypes through sensing of tissue distress signals^74^. Recent studies have identified discrete subsets/states of microglia that emerge during the progression of pathology in neurodegenerative disease models that in part involves repression of some constitutively-expressed homeostatic genes^75^. We observed marked suppression of the homeostatic genes *P2ry12* and *P2ry13* during the induction phase that was largely restored during resolution. This is similar to LPS-induced suppression of these genes in cultured microglia. The resolution cluster contained many other homeostatic genes (e.g. *Fcrls*,*Hexb*, *Olfml3*, *Tmem119*) and their selective induction during the Resolution phase alongside restored *P2ry12/13* expression is intriguing. The transcriptome data are derived from tissue homogenates therefore we cannot completely exclude that this pattern could reflect cell (e.g. microglial cell numbers). However, this is unlikely because there were negligible differences in microglial numbers quantified by flow cytometry during the progression of the inflammatory response. Moreover, the suppression of *P2ry12/13* at the same time as induction of large arrays of cytokine and other inflammatory gene families suggests phenotypic changes in microglia are a more likely explanation. Restoration of a homeostatic microglial phenotype may be an important element of inflammation resolution however if this is a largely end-stage effect or if the induction of homeostatic networks is actively involved in resolution cannot be concluded from the selected timepoints we studied. With more extensive temporal profiling and single cell transcriptomics it could be possible to identify if initial deviation from a homeostatic state is necessary for microglia to engage a reactive pro-resolution phenotype that subsequently returns towards a pre-inflamed steady-state profile. This question mirrors the current debate around the possible harms and benefits of microglia deviating from their homeostatic state during disease^55^ and will be important to address so that contexts where controlled manipulation of microglia to promote retention or deviation from steady-state will be beneficial can be determined.

In the present study, we have focussed on understanding key cellular and molecular changes in a model of self-limiting brain inflammation largely focussing on the early stage of resolution during which clearance of initial inflammatory cell infiltrates is prominent. Resolution will involve an array of coordinated processes and may involve more chronic changes that are important for long-term restoration of tissue homeostasis. We acknowledge these will be important to study and build upon the framework presented here. For example, and as implicated by our transcriptome profiling discussed above, understanding how the microglial population returns to true steady state in the later resolution phase will be important. In conclusion, we have shown how resolution of acute inflammation in the brain involves active induction of cellular and molecular events, have identified candidate gene networks that may regulate resolution-phase adaptations and propose the current model as a useful benchmark and framework for further understanding mechanisms and modifiers of inflammation resolution in the brain. The data also imply that inflammation-modifying strategies to treat CNS pathology are unlikely to be optimal if they are too indiscriminate and suppressive of adaptive changes integral to pro-resolution mechanisms.

## Supporting information

## Author contributions

BWM conceived study; BWM and CLD designed and performed experiments and analysed all data; AP contributed to analysis and presentation of transcriptomic data; all authors contributed to manuscript writing and editing.

## Acknowledgments

We thank staff in the Roslin Institute Biological Research Facility for animal husbandry and technical assistance and Dr Anna Raper for assistance with flow cytometry. We are grateful to Dr Daniel Anthony (University of Oxford) for kind provision of the SJC antibody. Microarrays were performed by Edinburgh Genomics (http://genomics.ed.ac.uk/).

